# Oscillatory population-level activity of dorsal raphe serotonergic neurons sculpts sleep structure

**DOI:** 10.1101/2021.11.19.469231

**Authors:** Tomonobu Kato, Yasue Mitsukura, Keitaro Yoshida, Masaru Mimura, Norio Takata, Kenji F. Tanaka

## Abstract

Dorsal raphe (DR) 5-HT neurons are involved in regulating sleep-wake transitions. Previous studies demonstrated that single-unit activity of DR 5-HT neurons is high during wakefulness, decreases during non–rapid eye movement (NREM) sleep, and ceases during rapid eye movement (REM) sleep. However, characteristics of the population-level activity of DR 5-HT neurons, which can influence the entire brain, are largely unknown. Here we measured population activities of 5-HT neurons in male and female mouse DR across the sleep-wake cycle by a ratiometric fiber photometry system. We found a slow oscillatory activity of compound intracellular Ca^2+^ signals during NREM sleep. The trough of concave 5-HT activity increased along with sleep progression, but the 5-HT activity level always returned to that seen in wake periods. When the trough reached the minimum level and remained there, REM sleep initiated. We also found a unique coupling of the oscillatory 5-HT activity and EEG power fluctuation, suggesting that EEG fluctuation is a proxy for 5-HT activity. Optogenetic activation of 5-HT neurons during NREM sleep triggered a high EMG power and induced wakefulness. Optogenetic inhibition induced REM sleep or sustained NREM with an EEG power increase and EEG fluctuation. These manipulations demonstrated a causal role of DR 5-HT neurons in sculpting sleep-wake structure. We also observed EEG fluctuations in human males during NREM sleep, implicating the existence of 5- HT oscillatory activity in humans. We propose that NREM sleep is not a monotonous state, but that it is dynamically regulated by the oscillatory population activity of DR 5- HT neurons.

**Significant statement:** Previous studies have demonstrated single-cell 5-HT neuronal activity across sleep- wake conditions; however, population-level activities of these neurons are largely unknown. We monitored dorsal raphe (DR) 5-HT population activity using a fiber photometry system in mice and demonstrated that activity was highest during wakefulness, and lowest during rapid eye movement (REM) sleep. Surprisingly, during non-REM (NREM) sleep, the 5-HT population activity decreased with an oscillatory pattern, coinciding with EEG fluctuations. We examined the causal role of these 5-HT neuron activities by optogenetics and found that DR 5-HT neurons sculpted sleep-wake conditions by influencing EEG and EMG patterns. We found similar EEG fluctuations in a human sleep EEG study, suggesting the presence of oscillatory 5-HT neuron activity during NREM across species.

## Introduction

The dorsal raphe (DR) nucleus in the hindbrain contains about one third of the brain’s 5-HT neurons (Müller et al., 2010). DR 5-HT neurons mainly innervate the forebrain, including the cortex and the striatum (Gaspar and Lillesaar, 2012). The roles of DR 5- HT neurons vary from regulating physical activities to regulating emotional states (Müller et al., 2010).

In regards to sleep-wake regulation by DR 5-HT neurons, studies employing DR 5-HT neuron ablation or other methods that induce a loss-of-function of these neurons demonstrated disturbances in sleep-wake structure. DR ablation reduced sleep in cats (Jouvet, 1968) and fish (Oikonomou et al., 2019). Administration of the irreversible inhibitor of tryptophan hydroxylase (Tph; the rate-limiting enzyme in 5-HT synthesis) reduced sleep in monkeys (Weitzman et al, 1968), cats (Koella et al, 1968), rats (Mouret et al.,1968; Torda, 1967), and fish (Oikonomou et al., 2019). Knockout of *Tph2* (the gene encoding the central nervous system isoform of Tph) reduced sleep (Siesta) in mice (Whitney et al., 2016) and fish (Oikonomou et al., 2019). All these long-term loss-of-function manipulations of the DR 5-HT neurons resulted in decreased sleep. In contrast, temporary cooling of the DR *induced* sleep in cats (Raymond et al., 1976), indicating an opposing outcome after acute loss-of-function of DR 5-HT neurons. Although the outcome of loss-of-function studies targeting DR 5-HT neurons are controversial, it is widely accepted that DR 5-HT neurons are causally involved in the sleep-wake structure.

Unlike the interventional studies modulating DR 5-HT neuronal activity, observational electrophysiological studies monitoring DR neuronal activities have reported consistent results (Mcginty & Harper, 1976; Lacher, 1985; Urbain et al., 2006; Sakai, 2011). Single-unit recording from the DR across the sleep-wake cycle revealed two major types of neurons. One type—the 5-HT neuron cell type—tonically fires during wakefulness, is less active during non–rapid eye movement (NREM) sleep, and is mostly silent during rapid eye movement (REM) sleep. The other cell type—non–5- HT neurons, such as dopaminergic or GABAergic neurons —does not modulate its firing rate across the sleep-wake cycle (Sakai, 2011). These pioneering studies encouraged us to monitor the population-level activity of DR 5-HT neurons across the sleep-wake cycle because the population 5-HT neuron activity, rather than individual neuron activity, could mediate a wide range of cortical activity patterns.

In this study, we sought to examine the dynamics of the population activity of DR 5-HT neurons during sleep in mice and to address how such dynamics were correlated to EEG and EMG changes. We addressed the causal relationship between DR 5-HT neuron activity and EEG/EMG changes across sleep stages using optogenetics. We further attempted to find similar EEG fluctuations in humans during NREM sleep.

## Materials and Methods

### Ethics statement

All animal procedures were conducted in accordance with the National Institutes of Health *Guide for the Care and Use of Laboratory Animals* and approved by the Keio University Animal Experiment Committee in compliance with the Keio University Institutional Animal Care and Use Committee (approval numbers: 12035 and 14027). Human experiments were approved by the Keio University Faculty of Science and Technology Bioethics Committee (approval ID: 31-7). This study was conducted following the principles of the Declaration of Helsinki. Informed consent was obtained from all participants.

### Animals

Experiments were conducted with 8-14–month-old male and female mice. All mice were maintained on a 12:12 h light/dark cycle (lights on at 08:00), and polysomnographic recordings were performed during the light phase. Tph2–yellow cameleon (YC) mice (*Tph2*-tTA::tetO-YC-nano50 double transgenic mice) were obtained by crossing tetO-YC-nano50 mice and *Tph2*-tTA mice (Miyazaki et al., 2014) and tetO-YC-nano50 mice (Kanemaru et al., 2014). Tph2-ChR2 mice (*Tph2*-tTA::tetO- ChR2(C128S)-EYFP double transgenic mice) were obtained by crossing *Tph2*-tTA mice and tetO-ChR2 mice (Tanaka et al., 2012). Tph2–Archaerhodopsin T (ArchT) mice (*Tph2*-tTA::tetO-ArchT-EGFP double transgenic mice) were obtained by crossing *Tph2*-tTA mice and tetO-ArchT-EGFP mice (Tsunematsu et al., 2013). All mouse lines were sourced from the RIKEN BioResource Center. The genetic background of all transgenic mice was mixed with C57BL6 and 129 SvEvTac. Genotyping for *Tph2*-tTA, tetO-YC-nano50, tetO-ChR2(C128S), and tetO-ArchT has been previously described (Tanaka et al., 2012; Tsunematsu et al., 2013; Miyazaki et al., 2014; Kenamaru et al., 2014).

### Surgical procedure

Surgeries were performed using a stereotaxic apparatus (SM-6M-HT, Narishige). Mice were anesthetized with a mixture of ketamine and xylazine (100c: g/kg and 10c:mg/kg, respectively). Body temperature during surgery was maintained at 37 ± 0.5°C using a heating pad (FHC-MO, Muromachi Kikai). An optic fiber for optogenetics or photometry was inserted into the DR at the following coordinates relative to bregma: AP, −4.3 mm; ML, 0.0 mm; and DV, 3.0 mm, all while tilted at 10° relative to the vertical axis (SM-15R, Narishige). The mice received permanent EEG and EMG electrode implants for polysomnography. Using a carbide cutter (drill size diameter: 0.8 mm), three pits were drilled into the skull, while avoiding penetration of the skull to prevent brain damage. Each implant had a 1.0-mm diameter stainless steel screw that served as an EEG electrode—one implant was placed over the right frontal cortical area (AP: +1.0 mm; ML: +1.5 mm) as a reference electrode and the other over the right parietal area (AP: +1 mm anterior to lambda; ML: +1.5 mm) as a signal electrode. Another electrode was placed over the right cerebellar cortex (AP: −1.0 mm posterior to lambda; ML: +1.5 mm) as a ground electrode. Two silver wires (AS633; Cooner Wire Company, USA) were placed bilaterally into the trapezius muscles and served as EMG electrodes. Finally, the electrode assembly and optical fiber cannula were anchored and fixed to the skull with Super-Bond (Sun Medical Co., Shiga, Japan).

### EEG/EMG recordings

The EEG/EMG signals were amplified (gain ×1000) and filtered (EEG: 1-100c:Hz, EMG: 10-100c:Hz) using a DC/AC differential amplifier (AM-3000, AM systems). The input was then received via an input module (NI-9215, National Instruments), digitized at a sampling rate of 1000 Hz by a data acquisition module (cDAQ-9174, National Instruments), and recorded by a custom-made LabVIEW program (National Instruments). We habituated the mice sufficiently, in other ward, REM sleep (see Vigilance State Assessment) was often observed, then started measurements.

Measurements were performed on 5 mice at 1-4 h/session, 1-2 sessions/mice (total 7 sessions).

### Mice Vigilance state assessment

EEG/EMG signals were analyzed using MATLAB (MathWorks, MA, USA). The power spectral data of the EEG were obtained using the multispectrogram method. A power spectral profile over a 1-50 Hz window was used for the analysis. We detected each sleep-wake state scored offline by characterizing 1-s epochs, as described in a previous study(Funato et al., 2016). A wake state was characterized by low-amplitude fast EEG and high-amplitude variable EMG. NREM sleep was characterized by high- amplitude delta (1-4 Hz) frequency EEG and a low-amplitude tonus EMG. REM sleep was staged based on theta (6-9 Hz)-dominant EEG and EMG atonia.

### Fiber photometry

The method for ratiometric fiber photometry has been described previously (Natsubori et al., 2017). An excitation light (435c:nm; silver light–emitting diode, Prizmatix) was reflected off a dichroic mirror (DM455CFP, Olympus), focused with a 2× objective lens (numerical aperture 0.39, Olympus) and coupled into an optical fiber (M79L01, Φ 400μm). The light-emitting diode power was <200c:μW at the fiber tip. The cyan and yellow fluorescence emitted by YC-nano50 was collected via an optical fiber cannula, divided by a dichroic mirror (DM515YFP, Olympus) into cyan (483/32 nm band path filters, Semrock) and yellow (542/27c:nm), and detected by a photomultiplier tube (H10722-210, Hamamatsu Photonics). The fluorescence signals were digitized using a data acquisition module (cDAQ-9174, National Instruments) and simultaneously recorded using a custom-made LabVIEW program (National Instruments). Signals were collected at a sampling frequency of 1000c:Hz.

### Optogenetic manipulation

An optical fiber (numerical aperture 0.39, Thorlabs) was inserted through the guide cannula. Blue (470c:nm) and yellow (575c: m) light was generated using a SPECTRA 2-LCR-XA light engine (Lumencor). The blue and yellow light power intensity at the tip of the optical fiber was 0.5-1c:mW and 6-8c:mW, respectively. During EEG and EMG monitoring, we illuminated ChR2-expressing mice during the wake, NREM, and REM periods. We illuminated Arch-T–expressing mice during the NREM period. For ChR2 activation, blue and yellow light (1c: and 5 s duration, respectively) were used to open and close the step-function type opsin ChR2(C128S) (Berndt et al., 2009). In the control trials, yellow light was used instead of blue light in Tph2-ChR2 mice. For ArchT activation, a 120-s duration of yellow (inhibition) light was used in Tph2-ArchT mice. In control trials, yellow light was used in wild-type mice.

### Histology

Mice were deeply anesthetized with ketamine (100c:mg/kg) and xylazine (10c:mg/kg) and perfused with 4% paraformaldehyde phosphate buffer solution. Brains were removed from the skull and postfixed in the same fixative overnight. Subsequently, the brains were cryoprotected in 20% sucrose overnight, frozen, and cut at a 25- μm thickness on a cryostat. Sections were mounted on silane-coated glass slides (Matsunami Glass). The sections were incubated overnight with anti-GFP antibodies (1:200, goat polyclonal, Rockland) at room temperature and then incubated with anti-goat IgG antibody conjugated to Alexa Fluor 488 (1:1000, Invitrogen) for 2c: at room temperature. Fluorescence images were obtained using an all-in-one microscope (BZ- X710, Keyence).

### Human subjects and polysomnography

We included 9 healthy male participants (aged 20-29 years [mean ± SD = 23.6 ± 0.22]). The exclusion criteria were (a) a history of neurological or psychiatric diseases and (b) alcohol or drug abuse. A total of 10 recording electrodes were prepared, including 4 EEG channels (C3A2, C4A1, O1A2, O2A1), 2 EOG channels (LOCA1, LOCA3), and 3 EMG channels (chin, both knees). All recordings were sampled at a rate of 200 Hz.

EEG recording were made using Ag/AgCl electrodes. Data were acquired with polysomnography equipment (Philips Healthcare, Alice PDx), and the sleep stages were judged by the American Academy of Sleep Medicine scoring rules.

### Data processing and analysis

All animals and trials were randomly assigned to an experimental condition. Experimenters were not blinded to the experimental conditions during data collection and analysis. Mice were excluded when the optical fiber position was not correctly targeted. Fiber photometry data were analyzed using custom-made programs in MATLAB. Yellow and cyan fluorescence were fitted using a binary exponential function to counteract the fading of fluorescent proteins and the fading of autofluorescence of optical fibers. We then used the YC ratio (*R*), which is the ratio of yellow to cyan fluorescence intensity, for calculating neural activity. We derived the value of the photometry signal (Δ*R/R_0_*) by calculating (*R* – *R*_0_) / *R*_0_, where *R* was the baseline fluorescence signal (signals in the wake state). For normalization of activity intensity, population 5-HT activities during the wake period were regarded as 0, and the REM period was regarded as −1. We defined the baseline at −0.1 to omit small fluctuations and/or baseline trends. We defined the single concave wave as the event that had a trough below −0.5. For normalization of the length of each epoch (one session of each sleep state), we normalized the length at 1000 (a.u.). Then, we calculated the maximum and minimum 5-HT activities for every 100 (a.u.). Data for all experiments were analyzed using parametric statistics: Student’s *t* test (Independent- samples *t*-test), paired *t* test, and one-way ANOVA followed by the Tukey-Kramer post hoc test, as well as repeated measures ANOVA. For mouse EEG, we set each EEG frequency band as follows: delta: 1-4 Hz; theta: 6-9 Hz; alpha; 9-12 Hz; and beta: 12-30 Hz (Choi et al., 2010). In humans, we set it as follows: delta: 1-3 Hz; theta: 4-7 Hz; alpha: 8-13 Hz; and beta: 15-28 Hz (Başar et al., 2013).

### Data availability

The datasets generated during and/or analyzed in the current study are available from the corresponding author on reasonable request.

## Results

### Dorsal raphe 5-HT neurons showed oscillatory population-level activity during

#### NREM sleep

To investigate population activity of 5-HT neurons in the DR during the sleep-wake cycle, we used a fiber photometry system and monitored intracellular calcium signals from 5-HT neurons in the DR of freely moving mice (**Fig. 1a**). We used transgenic mice expressing a FRET–based ratiometric Ca^2+^ indicator, YC-nano50 (Horikawa et al., 2010), in 5-HT neurons under the control of the *Tph2* promoter (Tph2-YC mice; **Fig. 1b, c**). The ratio of yellow to cyan fluorescence intensities (YC ratio) represents a compound Ca^2+^ activity of 5-HT neurons. Since fluorescence intensities of these two colors exhibit inversely proportional dynamics according to changes in Ca^2+^ concentration, the YC ratio is suited for detecting a decrease as well as an increase in Ca^2+^ concentration (Tsutsui-Kimura et al., 2017; Yoshida et al., 2019).

**Figure 1.**
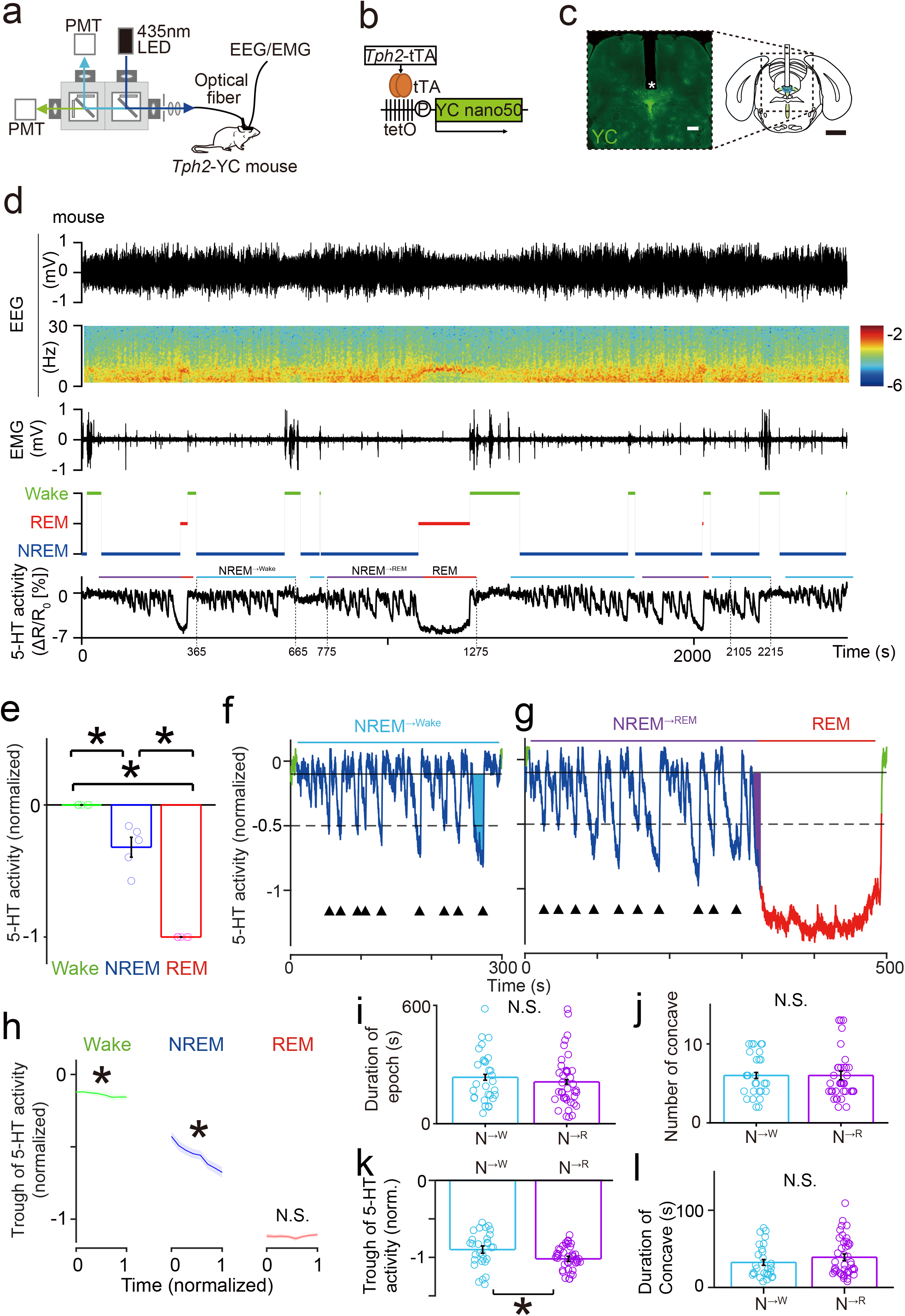
Dorsal raphe (DR) 5-HT neurons in mice showed oscillatory population activity during non–rapid eye movement (NREM) sleep. (a) Schematic illustration of a fiber photometry system for monitoring DR 5-HT neuron activity. EEG and EMG were recorded simultaneously. PMT: Photomultiplier tube. (b) Genetic construct of *Tph2*-tTA::tetO-YC-nano50 double transgenic mice. (c) The compound Ca^2+^ signal in the DR was monitored via an optic fiber (asterisk indicates the tip of the fiber). A fluorescence image shows YC-nano50 expression in the raphe nucleus. Scale bar, 1c:mm. (d) Representative examples of acquired data, including a trace of the EEG signal, a relative EEG power spectrum, a trace of the EMG signal, the sleep-wake states (wake: green; rapid eye movement [REM]: red; NREM: blue), and population Ca^2+^ dynamics of DR 5-HT neurons. Light blue lines indicate NREM^→Wake^ epochs, purple lines indicate NREM^→REM^ epochs, and red lines indicate REM epochs. Numbers along the bottom indicate time. (e) Normalized DR 5-HT neuron activities across the sleep-wake cycle. Averaged DR 5-HT neuron activity during NREM sleep was lower than during the wake stage, but higher than during REM sleep (one-way ANOVA followed by Tukey-Kramer post hoc test: F (2,12) = 138.4, wake vs NREM, p= 5.9×10^-4^; NREM vs REM, p= 3.3×10^-7^; wake vs REM, p= 4.0×10^-9^). \**p* < 0.05. (f and g). Representative population activity dynamics of DR 5-HT neurons during the NREM^→Wake^ epoch (f) and the NREM^→REM^ epoch (g). The solid and dashed lines show −0.1 and −0.5 of normalized activities, respectively. Waves that exceeded −0.5 are marked by black triangles. Time 0 in (f) corresponds to 365 seconds in (d), and time 0 in (g) corresponds to 775 seconds in (d). (h) Trough level of each wave gradually decreased over time during wake (F (9, 54) = 3.5, *p* = 1.6 × 10^-3^) and NREM (F (9, 54) = 43, *p* = 1.5×10^-21^, repeated measures ANOVA), but not during the REM (F (9, 54) = 0.68, *p* = 0.72). Shaded area indicates SEM. \**p* < 0.05. (i) Durations of the NREM^→Wake^ epoch (n = 29 epochs from 5 animals) and the NREM^→ REM^ epoch (n = 38 epochs from 5 animals) were comparable (*p* = 0.55, df = 65, t = 0.71, independent *t* test). Error bars indicate SEM. (j) Numbers of concave waves during NREM^→Wake^ and NREM^→REM^ were comparable (*p* = 1.0, df = 65, t = 0, independent *t* test). Error bars indicate SEM. (k) The trough of the last concave wave during NREM^→REM^ was lower than that during NREM^→Wake^ (*p* = 0.01, df = 65, t = 2.6, independent *t* test). Error bars indicate SEM. \**p* < 0.05. (l) Durations of the last concave wave in NREM^→REM^ and NREM^→Wake^ were comparable (*p* = 0.2, df = 65, t = -1.2, independent *t* test). Error bars indicate SEM.

We observed changes in the population activity of DR 5-HT neurons across the sleep-wake cycle (**Fig. 1d** bottom). Sleep-wake stages were identified with EEG and EMG measurements (**Fig. 1d**). During wake periods (green bars in the hypnogram; **Fig. 1d**), population activity of the DR 5-HT neurons showed small amplitude fluctuations. During NREM sleep (dark blue bars), the average population activity was lower than that during the wake period. During REM sleep (red bars), population activity of the DR 5-HT neurons was at a minimum (**Fig. 1e**; normalized 5-HT activities of 7 sessions from 5 mice, 1-4 h per sessions for total 17 h; mean 5-HT activities during wake and REM states were normalized to 0 and −1, respectively). These observations were consistent with previous findings obtained through electrophysiological studies examining single neuron activity (Jacobs et al., 1992).

We found oscillatory population activity of DR 5-HT neurons during NREM side and a REM period on the other (NREM^→REM^; **Fig. 1g**). Hereafter we defined both NREM epochs as a period including more than three concaves (downward Ca^2+^ waves) with a greater than −0.5 trough. Using these criteria, the duration of NREM^→Wake^ and NREM^→REM^ epochs was similar (**Fig. 1i**; mean ± SD: 196 ± 108 s and 177 ± 113 s, respectively; n = 29 and 38 epochs, respectively, from 5 animals). A small concave wave of 5-HT activity appeared during the first half to one-third of either NREM^→Wake^ or NREM^→REM^ epochs (**Fig. 1f, g**). The trough of the concave wave gradually declined during a NREM period (**Fig. 1h**). The 5-HT population activity of each concave wave returned from the trough to its baseline, which was similar to the activity level during a wake period. We then defined the concave 5-HT activity wave as the event that had a trough level below −0.5. There was no difference in the number of concave waves during an epoch of NREM^→Wake^ or NREM^→REM^ (mean ± SD: 6.0 ± 2.6 and 6.0 ± 3.2, respectively; **Fig. 1j**). Activity of 5-HT neurons of the last concave wave during NREM sleep (NREM^→REM^) did not return to its baseline, but decreased further to a lower level of a 5-HT activity, and then NREM switched to REM sleep. The mean trough level of the last concave wave during a NREM^→REM^ epoch (purple area in **Fig. 1f**) was slightly lower than that of a NREM^→Wake^ epoch (cyan area in **Fig. 1g**) (mean ± SD: −0.9 ± 0.2 vs −1.0 ± 0.2, *p* = 0.01, independent *t* test. **Fig. 1k**). There was no significant difference between the duration of the last concave wave in a NREM^→Wake^ epoch versus a NREM^→REM^ epoch (mean ± SD: 32 ± 21 and 39 ± 25 s, respectively; p = 0.2, independent *t* test **Fig. 1l**). Together, the lower trough level of the last concave activity, rather than the duration of an NREM epoch, the number of concave waves in NREM, or the duration of the last concave wave, was associated with a transition from NREM to REM.

### Low DR 5-HT neuron population activity was accompanied by an increase in wideband EEG power

Lowered 5-HT neuron activity may result in altered cortical EEG signals because DR 5- HT neurons innervate most cortical regions (Gaspar and Lillesaar, 2012). We noticed vertical stripes in the heatmap of the EEG spectrogram during NREM sleep (**Fig. 2a** top panel; see also **Fig. 1d** second panel), and thus we asked whether cortical EEG fluctuation was associated with the oscillatory 5-HT activity during NREM sleep. To study these repeated EEG activities, we normalized EEG power at frequencies from 10- 50 Hz at frequency intervals of 1 Hz (**Fig. 2a** second panel). The time course of total EEG power from 10-50 Hz exemplified the periodic increase and decrease in the wideband EEG power during NREM sleep (**Fig. 2a** third panel) and showed a roughly inverse correlation to 5-HT activity (**Fig. 2a** bottom panel).

**Figure 2.**
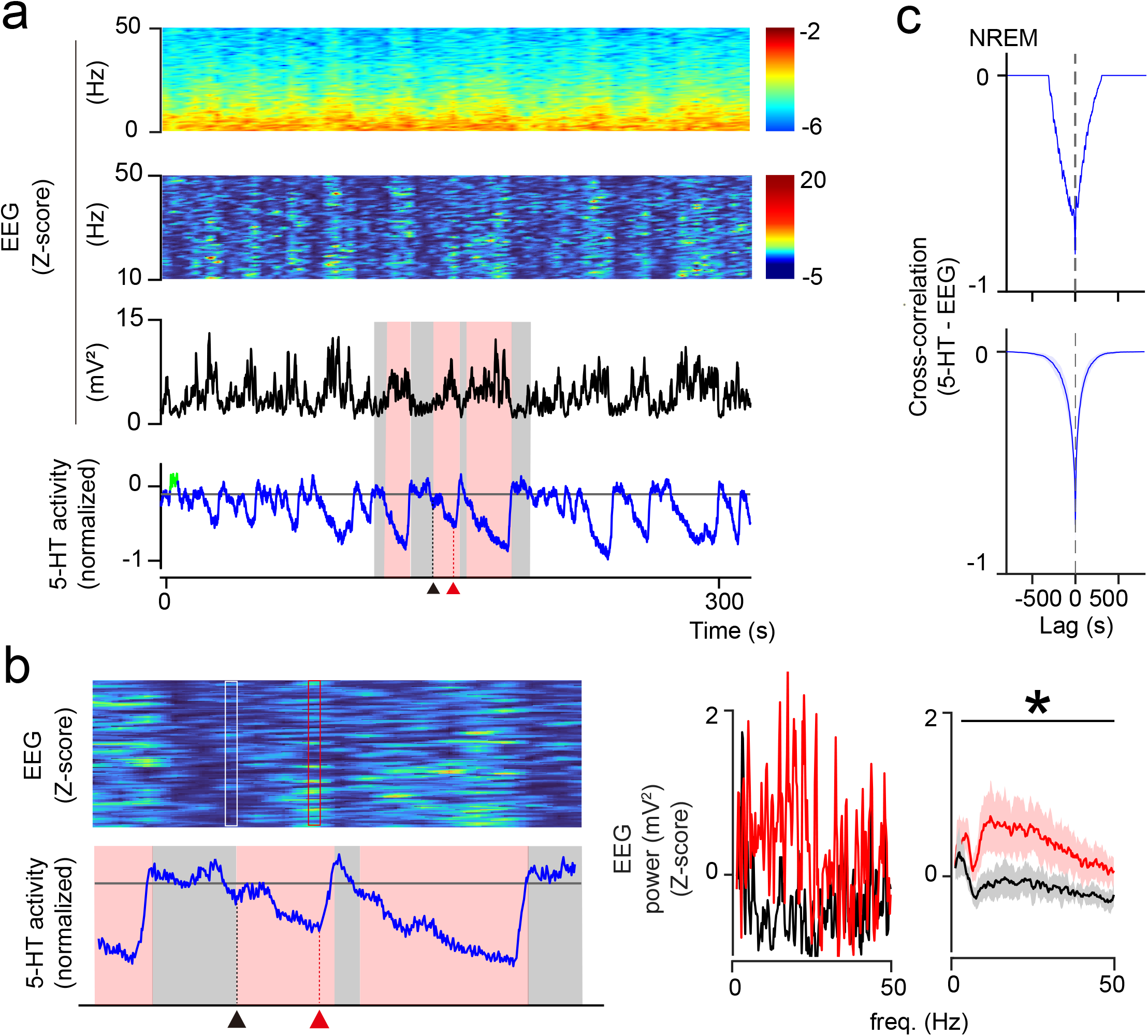
The increase in wideband EEG power repeatedly coincided with a decrease in dorsal raphe (DR) 5-HT neuron activity during non–rapid eye movement (NREM) sleep. (a) Temporal changes in the EEG power and the DR 5-HT neural activity. (top panel) The spectrum of relative EEG power. Periodic yellow-green stripes were observed. (second panel) Normalization of EEG power for every 1 Hz between 10-50 Hz. (third panel) The EEG power change. (bottom panel) DR 5-HT neural activity. Red shade indicates when 5-HT activity was below −0.1 and gray shade indicates when 5-HT activity was over −0.1; it is clear that low 5-HT neural activities coincided with a wideband EEG power increase and vice versa. In (a), 0 s corresponds to 775 s in Fig. 1d. Black arrowheads indicate the transition from gray to red shade. Red arrowheads indicate the trough in red shade. (b) (left panel) EEG data extraction ranges: the last 4 s of the basal 5-HT activity period (white box) and the 4 s before the trough of 5-HT activity (red box). (middle panel) individual data from the left panel. Red and black lines indicate powers in the white and red boxes, respectively. (right panel) population data (n = 5 mice, 7 sessions. Paired *t* test with a Bonferroni correction for every 1 Hz; black bar indicates \**p* < 0.01). The shaded area indicates SEM. (c) Cross-correlation between EEG power (10-50 Hz) and 5-HT activity during an NREM state. (top panel) Data from panel a. (bottom) Population data (n = 5). The shaded area indicates SEM.

We classified 5-HT activity during the NREM period into two categories: 1) basal activity when normalized 5-HT activity was at −0.1 or above (gray area in **Fig. 2a** (bottom) and **2b** (left)) a low activity state when the normalized 5-HT activity was less than −0.1 (red area in **Fig. 2a** (bottom) and **2b** (left)). We found a correspondence between the lowered activity of 5-HT neurons and a increase in the wideband EEG power (red area of EEG and 5-HT activities in **Fig. 2a**). To quantify the relationship, we extracted the normalized EEG power from the last 4 s of the basal period of 5-HT activity or from the 4 s before the trough of 5-HT activity, and compared the two EEG powers (**Fig. 2b**; n = 5 mice, 7 sessions, total 385 pairs). The EEG power during NREM at the point of low 5-HT activity was significantly larger than EEG power during basal 5-HT activity (**Fig. 2b**), confirming the augmentation of the EEG power during NREM with a lowered activity of 5-HT neurons. We found a weak, but significant, negative correlation between the trough level of the concave 5-HT activity and the magnitude of the EEG power across frequencies from 10-50 Hz (R = -0.13, p = 9.2×10^-3^, df = 768, n = 7 sessions from 5 mice, total 385 points). Next, we calculated a cross-correlation between the EEG power and the 5-HT activity during NREM sleep to investigate the temporal relationship between these parameters. The cross-correlation had a negative peak of −0.8 ± 0.1, with a lag of −0.3 ± 2.0 × 10^-2^ s and a full-width at half maximum of 59 ± 23 s (**Fig. 2c**; n = 5 mice 7 sessions). The lag was negligible because the time resolution of YC-nano50 for measurement of 5-HT activity was second. In summary, we found a transient and repetitive wideband EEG power increase that was associated with lowered 5-HT activity during NREM sleep.

### Activation of the 5-HT neurons induced wakefulness

In addition to the discovery of transient and repetitive increases in EEG power, we found occasional EMG amplitude increases during NREM sleep. It is well recognized that EMG amplitude increases appear at the transition from sleep to wake (**Fig. 1d**) (Funato et al., 2016). Indeed, we found an increase in population DR 5-HT neuron activity at the transition from sleep (both NREM and REM) to wake (**Fig. 1d**).

Therefore, it is possible that the increase in EMG amplitude during NREM sleep is associated with an increase in 5-HT neuron activity. We aligned the 5-HT neuron activity and EEG power during an NREM state and found the that 5-HT neuron activity was elevated concurrently with an EMG power increase (**Fig. 3a**). Cross-correlation analysis between myoelectricity and the population 5-HT neuron activity demonstrated that the time when 5-HT neuron activity returned to the wake level preceded the appearance of myoelectric activity by 0.6 seconds (**Fig. 3b**), suggesting that DR 5-HT neuron activity induced an EMG amplitude increase during NREM sleep.

**Figure 3.**
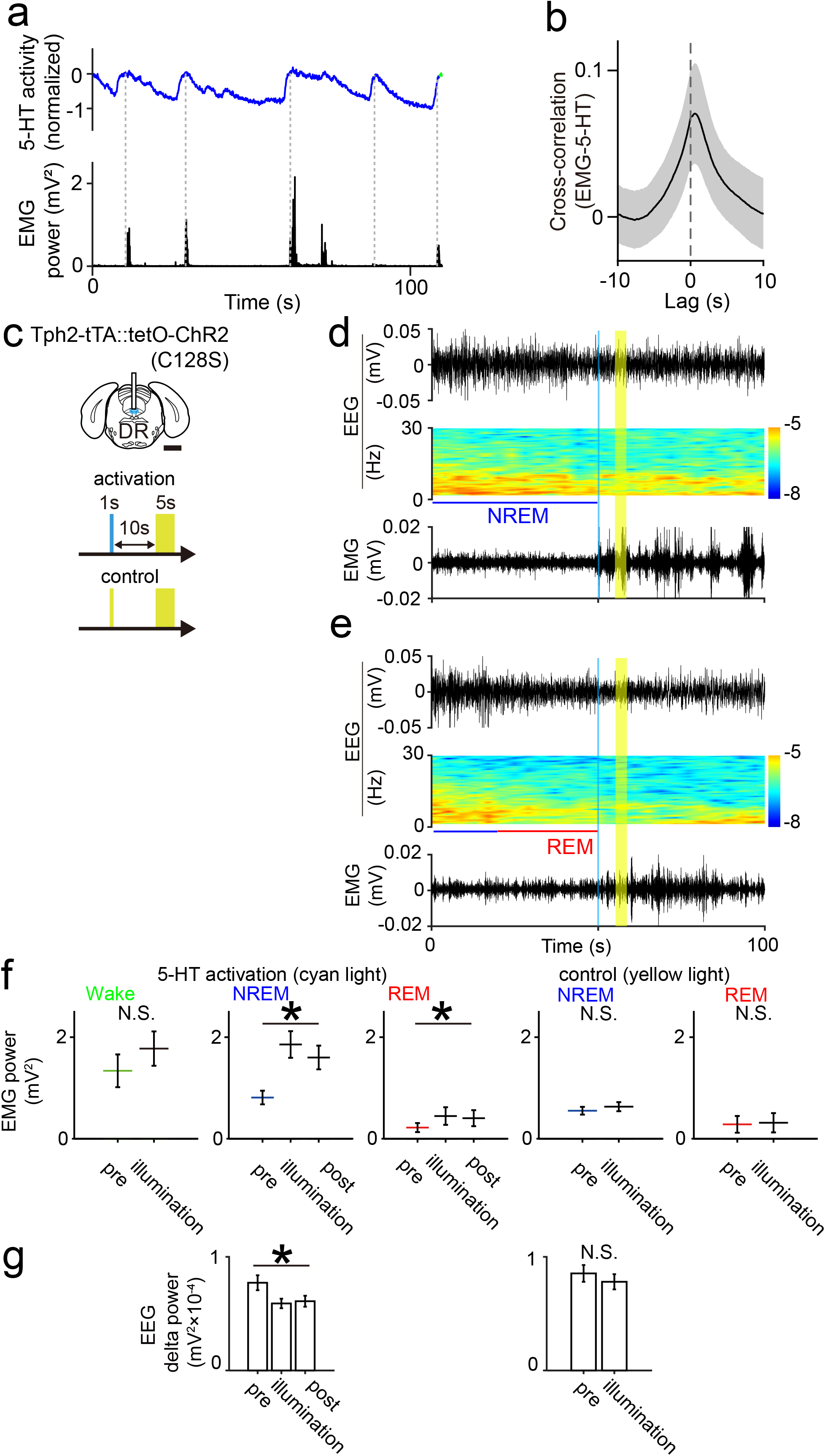
Activation of the dorsal raphe (DR) 5-HT neurons induced an EMG amplitude increase and wakefulness. (a) Temporal changes in population DR 5-HT neural activity (top) and EMG power during non–rapid eye movement (NREM) sleep (bottom). Vertical dotted lines indicate the peak timepoints when the declined DR 5-HT neuron activities returned to baseline. In panel a, 0 s corresponds to 2105 s in Fig. 1d. (b) Cross-correlation between DR 5-HT neural activity and EMG power during NREM sleep. The peak of EMG power followed the rise in DR 5-HT neuron activity with a 0.6-s delay. (c) Schematic illustration of optogenetic activation of DR 5-HT neurons in a Tph2- ChR2(C128S) mouse. Scale bar, 1c: m. Blue and yellow indicates illumination times. (d and e) EEG, EEG power spectrum, EMG, before and after optogenetic activation during NREM sleep (d) and rapid eye movement (REM) sleep (e), respectively. The vertical yellow lines indicate the timings for illumination. (f) Mean EMG power for the 10 s before, during, and after optogenetic activation. An EMG power increase was triggered by optogenetic activation and sustained afterwards (NREM: F(1, 2) = 20, p = 4.6×10^-2^, REM: F(1, 2) = 26, p = 3.6×10^-2^; repeated measures ANOVA). DR 5-HT optogenetic activation during a wake period did not alter EMG power (p = 0.22, df = 14, t = -1.3, paired *t* test). Yellow light illumination did not alter EMG power (NREM: p = 0.09, df = 32, t = -1.8, REM: p = 0.61, df = 5, t = -0.5; paired *t* test). \**p* < 0.05. Error bar shows SEM. (g) Mean EEG delta power for the 10 s before, during, and after optogenetic activation with cyan light during NREM. The significant delta power decline was induced and sustained (F(1, 2) = 129, p = 7.6×10^-3^, repeated measures ANOVA). Yellow light illumination did not alter EEG delta power (p = 0.18, df = 35, t = 1.4, paired *t* test).

To determine the causal relationship between DR 5-HT neuron activity and the EMG amplitude increase during NREM sleep, we used transgenic mice in which only 5-HT neurons express the step-function type variant of ChR2 (Tph2-ChR2(C128S) (Miyazaki et al., 2014) (**Fig. 3c**) and artificially activated their DR 5-HT neurons for 10 s during NREM sleep. Mice received 1 s of blue light illumination to open ChR2, followed by 5 s of yellow light illumination to close ChR2, 10 s after the blue light illumination (**Fig. 3c** middle). Optogenetic activation immediately increased the EMG amplitude (**Fig. 3d**, **3f**; n = 4 mice, wake: 15 sessions, NREM: 38 sessions, REM: 10 sessions) and decreased the delta power of the EEG (**Fig. 3d, 3g**), indicating a transition to the wake state. The induced wake state persisted after optogenetic activation, lasting 58 ± 49 s (mean ± SD, n = 42 sessions). This duration was the same as that of the wake period seen in control conditions during the light phase of the circadian cycle (**Fig. 3- 1a**). In addition, the artificial activation of DR 5-HT neurons during REM sleep induced wakefulness (**Fig. 3e, 3f**). However, application of the control yellow light (**Fig. 3c**) to Tph2-ChR2(C128S) mice during their natural sleep state did not induce EEG or EMG changes (**Fig. 3f, 3g**; n = 4 mice, NREM: 33 sessions, REM: 6 sessions). Blue light illumination to wild-type mice did not induce the EEG or EMG changes during sleep (**Fig. 3-1c**; n = 3 mice, wake: 13 sessions, NREM: 22 sessions, REM: 17 sessions). Artificial activation of DR 5-HT neurons during the wake state did not change EEG or EMG amplitude (**Fig. 3f**, **Fig. 3-1b**). Collectively, these data indicated that the myoelectric activity seen in NREM sleep was triggered by the rise to a peak level of DR 5-HT neuron activity. In addition, artificial activation of DR 5-HT neurons was sufficient to switch from a sleep to a wake state.

### Inhibition of 5-HT neurons occasionally induced REM

At the transition from NREM to REM sleep, the oscillation of DR 5-HT neuron activity terminated and the lowered DR 5-HT neuron activity in NREM was shifted to that in REM sleep. We sought to determine whether a continuous inhibition of DR 5-HT neuron activity induced REM sleep. For this inhibition, we used transgenic mice (Tph2- ArchT mice) harboring an inhibitory opsin, ArchT, in 5-HT neurons (**Fig. 4a**). In order to choose the duration of illumination, we measured the latency from the starting timepoint of the last concave wave during a NREM^→REM^ epoch to the trough timepoint of the following REM (**Fig. 4b**) and found that it ranged from 22-94 s (mean ± SD: 48 ± 21 s, n = 32 transitions; **Fig. 4c**). We thus chose 120 s as the illumination duration, to allow for a margin of error.

**Figure 4.**
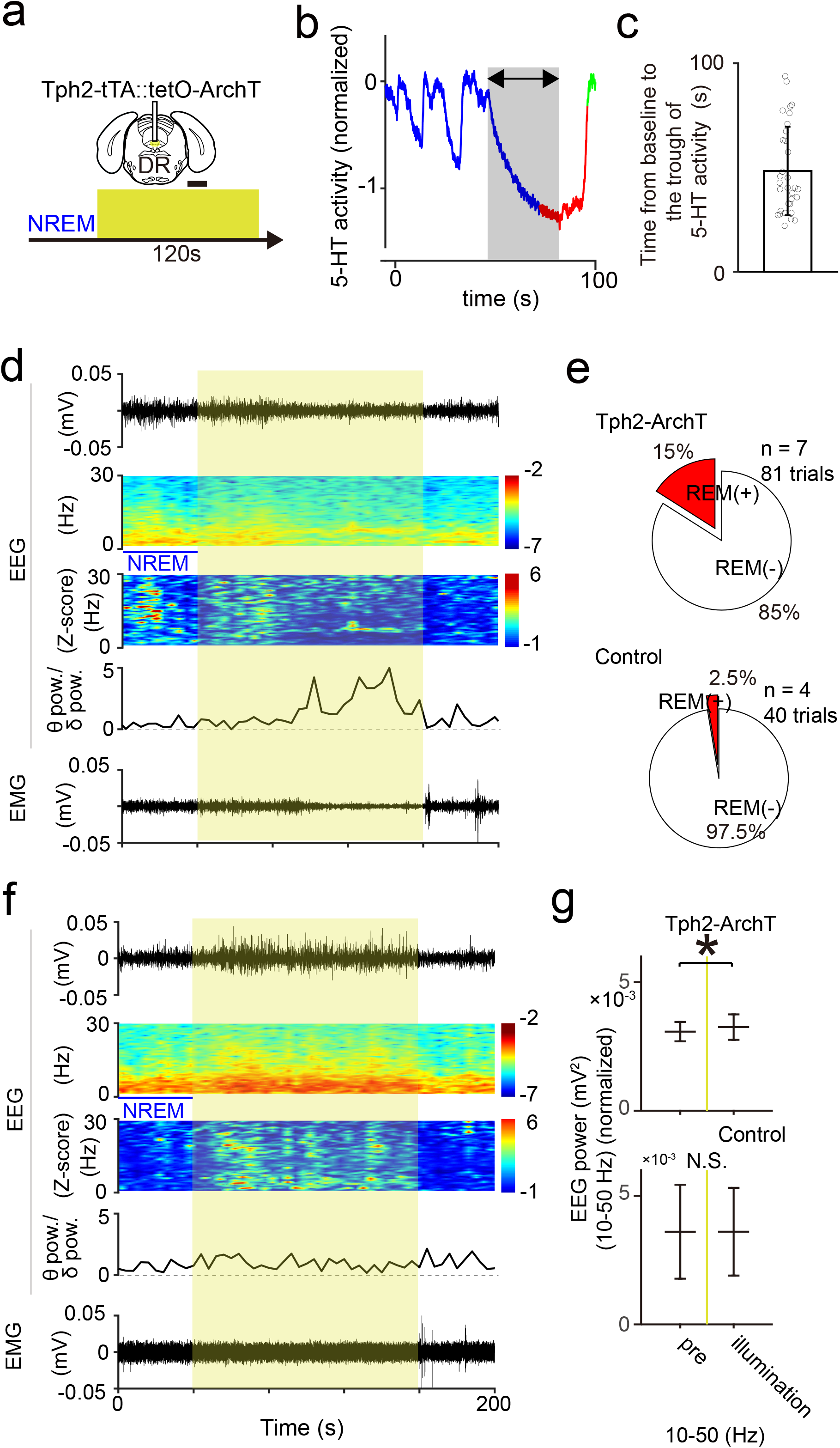
Inhibition of dorsal raphe (DR) 5-HT neuron activity increased the probability of rapid eye movement (REM) transition or sustained non–rapid eye movement (NREM). (a) Schematic illustration of optogenetic inhibition of DR 5-HT neurons in Tph2-ArchT mice. Scale bar, 1 mm. Yellow shade indicates light illumination. (b) Latency from the start time of the last concave wave of an NREM^→REM^ epoch to the time at the trough of the REM state that follows (gray). (c) Quantification of panel b (n = 32 transitions from NREM to REM). Data show mean ± SD. (d, f) Time courses of representative EEG, EEG power spectrum, normalized EEG power, ratio of theta power to delta power of EEG, and EMG with optogenetic inhibition (yellow), respectively from top to bottom. d) REM-induced trial, f) NREM- sustained trial. (e) Percentage of REM sleep–induced trials in Tph2-ArchT (7 mice, 81 sessions) and control mice (4 mice, 40 sessions). (g) Optogenetic inhibition induced a wideband EEG power increase (10-50 Hz: *p* = 0.03, df = 13, t = -2.5, paired t-test, \**p* < 0.05) and did not change in wild type mice (10-50 Hz: *p* = 0.99, df = 4, t = -5.0×10^-3^, paired t-test, \**p* < 0.05) in NREM sleep–sustained trials. Data show mean ± SD.

We applied optogenetic inhibition during a 3-h polysomnography recording during the light phase of the circadian cycle. We identified NREM sleep on-line and initiated 120 s of illumination at the first trial. We had an interval of at least 20 min before the next trial. As a result, we applied illumination 10 times on average during two recording sessions from each animal. In Tph2-ArchT mice (n = 7), out of 81 trials, 12 trials induced REM sleep (15%) (**Fig. 4d**). In controls (n=4), out of 40 trials, 1 trial induced REM sleep (2.5%) (**Fig. 4e**). In comparing these two stimulation conditions, we found that there is a higher probability of REM sleep induction by optogenetic inhibition versus control light stimulation (*p* = 0.04, Fisher’s exact test). Nonetheless, the REM sleep induction rate was low, and NREM sleep was sustained in most trials (79% vs 88%; Tph2-ArchT mice vs wild type mice). Of note, optogenetic inhibition after the third session did not increase the REM induction probability.

As we previously showed, oscillatory population DR 5-HT neuron activity corresponds to periodic EEG power fluctuations. Thus, we asked whether EEG power fluctuations could be perturbed by 120 s of optogenetic inhibition, a duration that should contain 3-4 concave waves of DR 5-HT neuron activity. Despite mice remaining in NREM sleep under illumination, we still found EEG power fluctuations (**Fig. 4f**), indicating two possibilities. The first is that our optogenetic inhibition was not enough to fully ablate the rise of DR 5-HT neuron activity during NREM. The other possibility is that oscillatory 5-HT neuron activity does not underlie EEG power fluctuation.

Artificial DR 5-HT silencing did not delete EEG fluctuation during NREM, however, we have always found a wideband EEG power increase during illumination (**Fig. 4f, 4g**), supporting the idea that DR 5-HT neuron inhibition during NREM is rather associated with an EEG power increase.

### The human brain also showed a transient and repetitive EEG power increase during NREM sleep

It is worth considering if the oscillation of 5-HT neuron activity during NREM sleep occurs in humans, even though humans have distinct sleep structures from mice. Namely, mice have monophasic sleep patterns, while humans have polyphasic ones. To pursue this question, we hypothesized that an infra-slow (<0.1 Hz) periodic increase of wideband EEG activity during NREM sleep could be a proxy for 5-HT activity in the human brain based on our demonstration in the mouse brain of the tight functional coupling of lowered 5-HT neuron activity and the heightened wideband EEG during NREM sleep. We obtained healthy human EEG/EMG/EOG data during a sleep-wake cycle (**Fig. 5a**). We observed a periodic transient increase in EEG power in the human brain during NREM sleep (**Fig. 5b**). To highlight these repeated convex activities of EEG, we normalized EEG power at frequencies from 5-30 Hz at frequency intervals of 1 Hz (**Fig. 5c**). Repeated convex EEG activities were evident during N2 of NREM sleep (black triangles in **Fig. 5c**). Note that in humans, NREM sleep is subdivided into three stages, N1, N2, and N3 (Carskadon and Dement, 2011).

**Figure 5.**
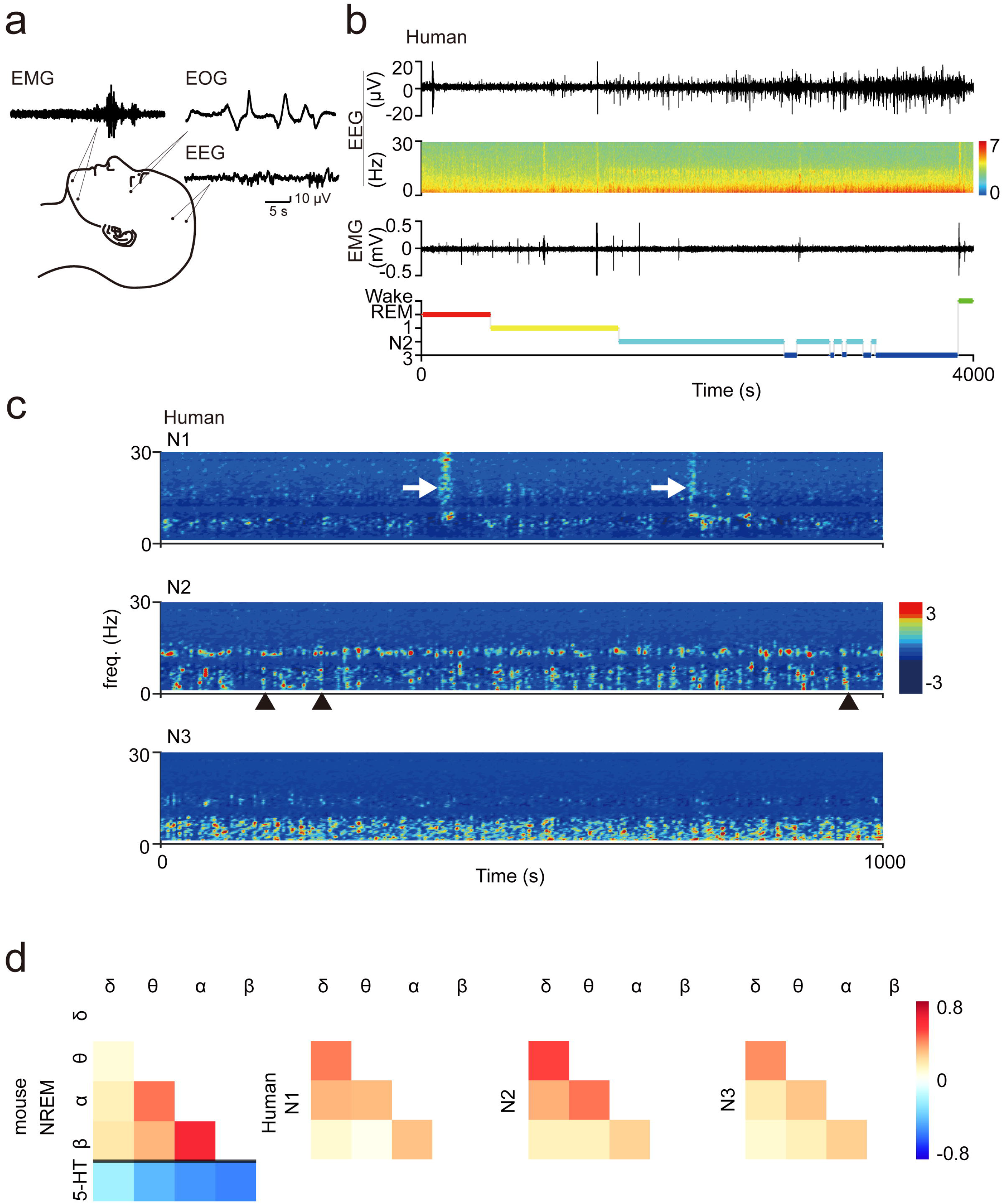
The human brain showed transient increases in EEG power during non– rapid eye movement (NREM). (a) Schematic diagram of polysomnographic recordings. (b) Representative EEG signal, EEG spectrogram, EMG signal, and hypnogram. (c) Z-scored EEG spectrograms in NREM stages 1-3 of a human subject. Black arrow heads in N2 show typical examples of transient increases in EEG power. White arrows in N1 show an EMG artifact. (d) Cross-correlation matrices of EEG signals at each frequency band and 5-HT activity during NREM sleep. In mice (leftmost panel), EEG signals at α vs β and α vs θ bands had moderate to strong positive cross-correlations. In humans, EEG signals at the same bands, namely α vs β and α vs θ, during all stages of NREM sleep showed weak to moderate positive cross-correlations. These results demonstrate that in both mice and humans there were higher cross-correlations among EEG signals in α, β, and θ frequencies, although the strongest correlation in human EEG was observed between δ vs θ bands. Note that the significant negative cross-correlations between 5-HT activity and EEG signal at α and β bands in mice may suggest that similar 5-HT dynamics occur in the human EEG.

To compare the frequency structure of the convex EEG activities of mice and humans, we calculated the cross-correlation coefficients between time-course of the EEG power at each frequency band during NREM sleep (**Fig. 5d**). Cross-correlation coefficients between EEG and 5-HT activities were also calculated for the mouse brain. In mice, strong positive cross-correlations were observed between EEG bands at α vs β (0.61 ± 0.03; mean ± SD) and α vs θ (0.45 ± 0.05; 1-2 sessions of 1-4 h recording from 5 mice each). The lowered 5-HT activity showed a strong negative cross-correlation to α and β bands of EEG (**Fig. 5d**, lower left section in the leftmost panel). These data demonstrated a relationship between EEG activities at each frequency band during a transient lowered 5-HT activity in the mouse brain. In humans, we observed strong positive cross-correlations between the same EEG bands at α vs β (N1: 0.28 ±0.14; N2: 0.25 ± 0.19; N3: 0.25 ± 0.09) and α vs θ (N1: 0.30 ± 0.05; N2: 0.43 ± 0.15; N3: 0.28 ± 0.09) during all stages of NREM sleep, although the strongest cross-correlation coefficient was found between the δ and θ bands (N1: 0.41 ± 0.13; N2: 0.51 ± 0.14; N3: 0.38 ± 0.07; **Fig. 5d** upper right; a single 8.5-h recording session of 5 human subjects). These data imply the possibility of oscillatory population activity of DR 5-HT neurons in the human brain.

## Discussion

The population 5-HT activity was high during wake, intermediate during NREM sleep, and low during REM sleep in average. We found a slow oscillatory population activity of 5-HT neurons (∼0.03 Hz) during NREM sleep. Oscillatory changes of population 5- HT activities coincided with dynamics of EEG power fluctuations in an anti-parallel manner.

Other groups have also described population-level 5-HT neuron activities by fiber photometry during sleep. Monitoring of DR 5-HT neuron activity using GCaMP6 revealed oscillatory patterns during NREM sleep (Oikonomou et al., 2019). The duration of the waves was similar to our findings, however, the signal did not return to the level seen in wakefulness periods. This difference may be due to the nature of the Ca^2+^ sensors used: YC is useful for detecting downward Ca^2+^ changes, while GCaMP was developed to efficiently detect spikes in Ca^2+^. Further study is needed to address this difference in dynamics using a distinct methodology. For example, the GPCR activation-based (GRAB)_5-HT_ sensor, a probe for extracellular 5-HT, would be ideal. This was recently examined in a study monitoring extracellular 5-HT levels in the basal forebrain, which revealed similar oscillatory dynamics during NREM sleep (Wan et al., 2021). Similar to the GCaMP6 study, though the released 5-HT levels fluctuated, the extracellular 5-HT levels did not return to the levels seen at periods of wakefulness.

With recent developments in GRAB_5-HT_ sensor technology, an improved version of it has been reported, having a similar Ca^2+^ detection pattern as that observed with YC (Japan Neuroscience meeting 2021, symposium); comprehensive results of this new technology are awaited.

In our photometry setup, we believe we captured a sufficient spatial range across the entire DR and were able to detect YC signals in 5-HT neurons in an unbiased manner. We used an optic fiber with a 400-µm diameter and a 0.39 numerical aperture. According to our previous estimation (Natsubori et al., 2017), this system could detect signals up to 700 µm beneath the tip of fiber. As a result, the shape of the range would be a conical frustrum, with 200- and 470- µm radiuses and a 700-µm height, suggesting that we covered most of the DR and did not detect signals from the median raphe (MR). A recent single-cell RNAseq study in DR/MR 5-HT neurons identified 6 clusters (Ren et al., 2019). Each cluster was located with some spatial bias in DR, but the range defined by the conical frustrum would roughly cover all clusters. Further, previous single-unit recordings from DR 5-HT neurons revealed four types of wake-activating neurons (that is, sleep-inhibiting neurons) and demonstrated their locations (Sakai, 2011). The location of each type varied, but the detection range of our photometry setup would cover most of DR neurons recorded. Together, we assume that we monitored most of DR 5-HT neurons in an unbiased manner in our photometry setup, including all the major clusters identified by single-cell RNAseq and the major cell types identified by single-cell recording. In fact, the patterns of DR 5-HT neuron population activities were indistinguishable between animals, supporting that the same population was targeted across animals in our study.

The population activity of DR 5-HT neurons was lowest during REM sleep. Does this mean that none of the DR 5-HT neurons fire during REM sleep? We believe this is not the case, since we did not observe the lowest level when REM sleep first began (**Fig. 1g, 4b**). It was only after a substantial delay that the population activity level reached its lowest, and still, it exhibited minor fluctuations afterward. To reconcile this decline at the initial phase of REM sleep, we reanalyzed a previous report (Sakai, 2011), in which the author monitored a total 229 DR 5-HT neuron activities by a single unit recording and described varied firing patterns during sleep. The author identified 195 wake-activating 5-HT neurons (4-5 Hz firing at wake periods), 9 wake/REM- activating probable 5-HT neurons (4-5 Hz firing at wake periods), and 25 sleep- activating probable 5-HT neurons (<0.5 Hz firing at wake periods). The author further classified wake-activating 5-HT neurons into 4 subtypes. Type I and II cells (n=115, 50%) completely ceased to fire at the REM sleep stage and type III and IV cells (n=80, 35%) fired at the beginning of REM sleep (<1 Hz) and decreased their firing rates (<0.5 Hz) afterwards. Wake/REM-activating neurons (4%) increased their firing rates from 2 Hz at the initial phase of REM to 4 Hz during REM sleep. Sleep-activating neurons (11%) fired at 2 Hz in both NREM and REM sleep. Since the population 5-HT neuron activity level at the beginning of REM sleep was higher than the level at the nadir during REM sleep, the decrease in firing rate in wake-activating type III and IV cells was attributable to the decline of population 5-HT neuron activity after the transition to REM. During REM sleep, wake/REM-activating neurons and sleep-activating neurons (total 15%) tonically fire at 2-4 Hz and these neuron activities may induce fluctuations during REM sleep.

REM sleep induction by optogenetic inhibition of DR 5-HT neurons is controversial. We were only able to induce REM sleep at the first and second sessions of illumination of ArchT. Illumination after the third session failed to induce REM sleep. These data suggested that silencing of DR 5-HT neurons facilitates the induction of REM sleep, but the effect of artificial silencing may be canceled adaptively. Although we induced REM sleep by optogenetic inhibition, the success rate was at most 15%, suggesting that another mechanism was required to induce REM sleep. In the future, it may worth trying to inhibit the locus coeruleus noradrenergic neurons because they fire at 1-3 Hz during the wake period, have reduced activity during NREM sleep, and cease firing during REM sleep (Aston-Jones and Bloom, 1981). Further studies are required to explore the conditions required to induce REM sleep efficiently.

Direct measurement of 5-HT dynamics in the human brain is difficult, though there are a few studies that have succeeded in detecting 5-HT fluctuations using invasive methods such as microdialysis or fast-scan cyclic voltammetry (Suominen et al., 2013; Silberbauer et al., 2019; Bang et al., 2020). Based on data in the present study, we propose a noninvasive strategy to infer 5-HT dynamics during NREM sleep in the human brain using EEG measurement: transient and repetitive EEG power surges, especially during NREM stage 2, may reflect decreased population 5-HT activities. In support of this idea, a relationship between EEG and 5-HT dynamics has been reported with pharmacological perturbations. Administration of selective serotonin reuptake inhibitors to healthy subjects reduced the total power of EEG (Dumont et al., 2005). In addition, a specific antagonist for serotonin 5-HT_2_ receptors prolonged the duration of slow-wave sleep and enhanced EEG power at delta and theta frequencies (Dijk et al., 1989).

In conclusion, the population activity of DR 5-HT neurons fluctuated during NREM sleep in mice. The temporal association between population DR 5-HT neuron activity and wideband EEG power and the outcomes of optogenetic manipulation of DR 5-HT neuron indicate that mouse NREM sleep is not a monotonous but a dynamic state sculpted by oscillatory population activity of DR 5-HT neurons.

---

**Figure 3-1.**
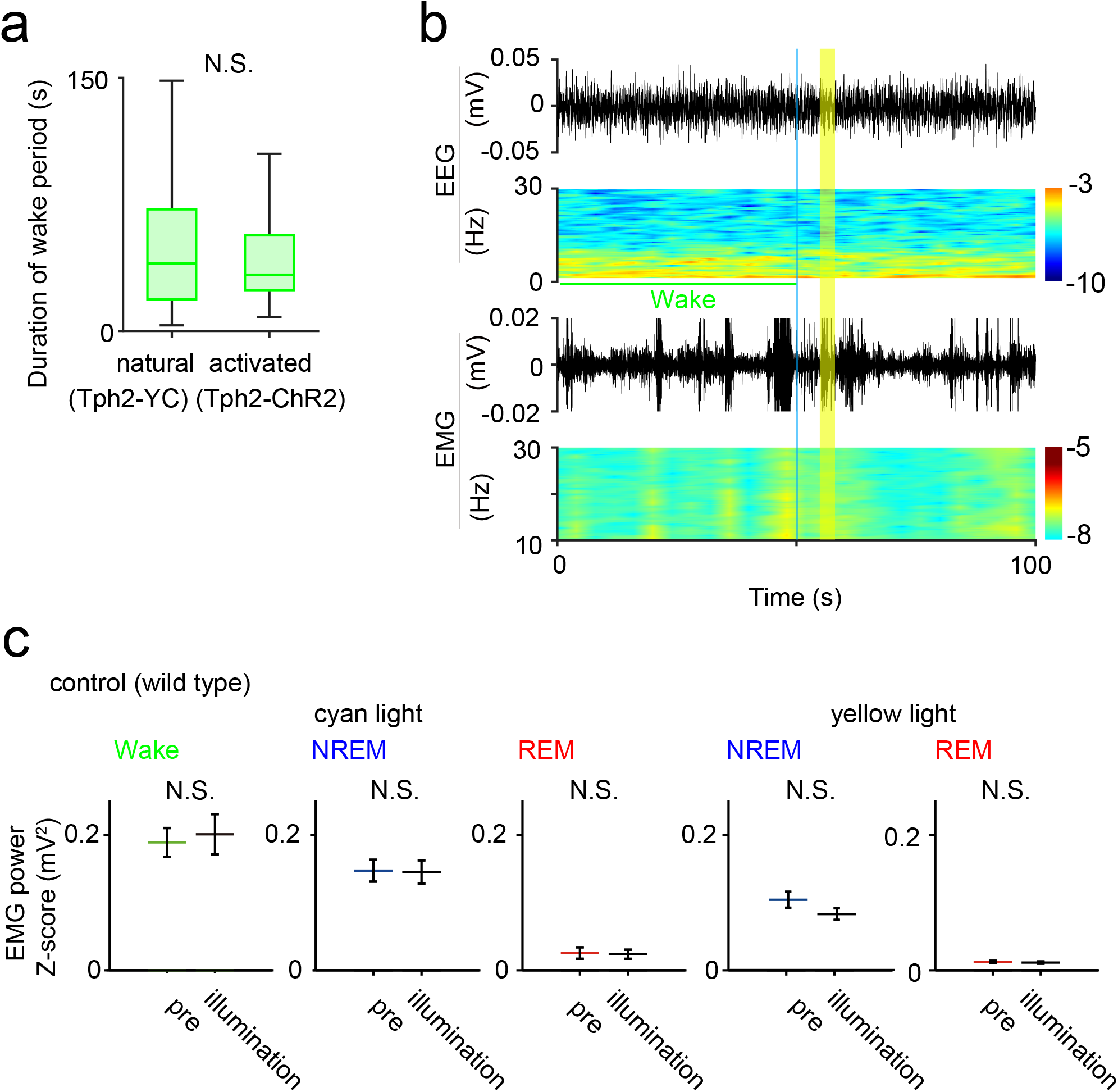
Light illumination of control mice did not change EMG power, related to Figure 3. (a) Duration of a wake period in Tph2-YC mice (left; n = 3, 1-2 sessions per mouse, 112 periods) and duration of the induced wake-like period after light illumination in Tph2- ChR2 mice (right; n = 3, 3 sessions per mouse, 42 periods). In the box plots, the central mark indicates the median, and the bottom and top edges of the box indicate the 25th and 75th percentiles, respectively. Whiskers denote the range. (b) Representative figures of the EEG, relative EEG power, EMG, and relative EMG power. Images represent data from Tph2-ChR2 mice that received cyan light illumination (optogenetic activation) during the wake period. Blue shade indicates 1 s of blue illumination and yellow shade indicates 5 s of yellow illumination. (c) Mean EMG power as measured 1 s before and after cyan light illumination (three left panels) or 1 s before and after yellow light illumination (two right panels) in control mice (n = 3, 3 sessions per mice). EMG power during the wake period (left side, left panel; total 13 illumination), during the non–rapid eye movement (NREM) period (left side, middle panel; total 22 illumination), and during the rapid eye movement (REM) period (left side, right panel; total 17 illumination), as well as EMG power following yellow light illumination as a control during NREM period (right side, left panel; total 24 illumination) and during the REM period (right side, left panel; total 13 illumination). There were no significant differences between pre and during illumination (cyan light: wake, p = 0.67, df = 12, t = -0.4; NREM, p = 0.89, df = 21, t = 0.15; REM, p = 0.41, df = 16, t = 0.85; yellow light: NREM, p = 0.06, df = 23, t = 2.0; REM, p = 0.21, df = 12, t = 1.3; paired *t* test).

## Notes

### Competing Interest Statement

The authors have declared no competing interest.

